# Transcriptomic analysis of the C3-CAM transition in Cistanthe longiscapa, a drought tolerant plant in the Atacama Desert

**DOI:** 10.1101/2022.03.16.484649

**Authors:** P. G. Ossa, A. A. Moreno, D. Orellana, M. Toro, T. Carrasco-Valenzuela, A. Riveros, C. C. Meneses, R. Nilo-Poyanco, A. Orellana

## Abstract

One of the most outstanding plant species during the blooming of the Atacama Desert is the annual plant *Cistanthe longiscapa*. This plant can perform CAM photosynthesis, but the ecophysiological and molecular mechanisms that this plant uses to withstand the extreme conditions it inhabits in the field are unknown.

Morphological and ecophysiological traits were studied and leaf samples at dawn/dusk times were collected from three sites distributed across an increasing south to north arid gradient, to evaluate CAM expression and transcriptomic differences, and search for links between photosynthetic path and abiotic response.

Plants from the different sites presented significant differences in nocturnal leaf acid accumulation, isotopic carbon ratio (δ^13^C), succulence and other four traits that clearly indicated a spectrum of CAM photosynthesis intensity that correlated with aridity intensity. The differential gene expression analysis among Dawn vs Dusk between sampling sites showed higher gene expression in the arid northern site (3991 v/s 2293) with activation of regulatory processes associated with abscisic acid and circadian rhythm.

The analysis highlights clear ecophysiological differences and the requirement of a strong rewiring of the gene expression to allow a transition from a weak into a strong CAM in *C. longiscapa*.

## Introduction

In the upcoming years, it is expected that temperatures and drought periods will intensify due to climate change, affecting biodiversity at all levels (Feng and Fu, 2013; Schlaepfer *et al*., 2017), with terrestrial plants particularly exposed to these stressful environmental factors (Feng and Fu, 2013; Parmesan and Hanley, 2015; Schlaepfer *et al*., 2017). Plants can sense environmental changes rapidly and evolve morphological, physiological, and molecular adaptations to face them (Dussarrat *et al*., 2018). However, the mechanisms underlying these responses are still unknown, making it difficult to build any prediction model on how plants will face the aforementioned challenges. In times of a fast-growing human population (Gerland *et al*., 2014), uncovering these mechanisms is essential for a sustainable agricultural development, and a relevant input to engineer more resistant crops for a warmer and drier world (Borland *et al*., 2014; Borland *et al*., 2015; DePaoli *et al*., 2014; Yang *et al*., 2015).

Photosynthesis is an essential plant metabolic process, where atmospheric CO_2_ is fixed into carbohydrates in a carbon cycle by RUBISCO, using the sun’s energy and water. However, this process is affected by drought and elevated temperatures (Perdomo *et al*., 2017). Under arid conditions, plants face a limited water income, which is countered by an active reduction of stomata opening. This strategy enables the plant to lose less water through transpiration, a trade-off that can be costly by limiting CO_2_ supply to photosynthesis (Bräutigam *et al*., 2017), and affecting the affinity of RUBISCO for CO_2_ instead of O_2_. To deal with this problem, some plants have evolved, together with adaptive morphological traits, CO_2_-concentrating mechanisms that can be coupled to carbon fixation in the Calvin-Benson cycle (Raven and Beardall, 2016; Raven *et al*., 2017). Crassulacean acid metabolism (CAM) is a CO_2_-concentrating mechanism remarkable for its high water-use efficiency (WUE) relative to C3 and C4 photosynthesis (Borland *et al*., 2009; Cushman *et al*., 2015). To reduce water loss, CAM plants fix CO_2_ during the night by the action of the enzyme phosphoenolpyruvate carboxylase (PEPC), generating organic acids (mainly malate) that are stored in the vacuole. During the day, the night accumulated organic acids are decarboxylated and CO_2_ is concentrated around RUBISCO to be fixed, entering the Calvin-Benson cycle (C3 photosynthesis) to form carbohydrates, also reducing photorespiration, which can have a detrimental effect over photosynthesis (Pereira *et al*., 2021). Given this temporal separation, CAM plants can achieve high WUE compared to C3 and C4 plants (Wai *et al*., 2019).

CAM is a remarkable example of convergent evolution of a complex trait that has evolved independently multiple times in about 6% of species and at least 37 plant families (Winter *et al*., 2020). Different types of CAM have been described (Messerschmid *et al*., 2021; Winter and Smith, 2022), for instance, in obligate CAM plants, like most cacti, carbon is fixed constitutively at night in contrast to facultative CAM plants, where nocturnal fixation is induced by drought, salt stress or ontogeny (Winter K., 2019; Schiller and Bräutigam, 2021). In CAM cycling plants, carbon is fixed nocturnally from recycled respiratory CO_2_, and during the day with C3 photosynthesis (Schiller and Bräutigam, 2021). CAM idling plants also perform CAM cycling, but stomata are always closed (Sipes and Ting, 1985). CAM species can also be classified as strong (CO_2_ night uptake >70%) or weak CAM (CO_2_ night uptake < 33%, Pereira *et al*., 2021). Plants performing weak CAM supplement the low-level nocturnal CO_2_ fixation with high daytime C3 photosynthesis carbon fixation (Winter K., 2019, Schiller and Bräutigam, 2021). These weak CAM states can be considered as intermediate steps within the evolutionary spectrum from C3 to CAM, or as the final evolutionary state for lineages where it might be advantageous (Hancock *et al*., 2019; Heyduk *et al*., 2019). On the other hand, depending on the route that takes malate decarboxylation during the day, two different CAM pathways can be distinguished (Holtum *et al*., 2005, Shameer *et al*., 2018): one is the malic enzyme route (CAM-ME), where malate is decarboxylated either in the cytosol by NADP-dependent malic enzyme (NADP-ME) or in the mitochondria by NAD-dependent malic enzyme (NAD-ME). These decarboxylations produce pyruvate and CO_2_, where pyruvate is transported to the chloroplast and converted to PEP by the chloroplastic PEP dikinase (PPDK). The other pathway is the PEP carboxykinase (PEPCK) route (CAM-PEPCK), where malate is decarboxylated by PEPCK producing PEP in the cytosol, so these species do not rely on PPDK. In addition, based on the transitory sugar accumulated, there are species where PEP is transiently stored as starch in the chloroplast, like in *Kalanchoe* or *Mesembryanthemum crystallinum;* and others where it is stored in the vacuole as soluble saccharides like sucrose, fructose or glucose (Holtum *et al*., 2005; Borland et al., 2016). These pathways diversity probably stems from the multiple evolutionary origins of CAM (Niechayev *et al*., 2019).

The evolutionary basis of CAM and its plasticity is sustained in the fact that all CAM genes exist in C3 species, with genetic changes centered on regulation, timing or abundance of transcripts, gene duplications and neofunctionalization (Heyduk *et al*., 2018, 2019). Changes in transcript abundance and regulation related to circadian rhythm have been proposed as the main source of changes related to CAM (Wai *et al*., 2019). Recently, changes in amino acid biosynthesis flux have also been suggested as a possible mechanism towards developing CAM photosynthesis (Bräutigam *et al*., 2017). In addition, genome-scale analyses of constitutive CAM plants suggest that time of day networks are phased to the evening compared to C3, whereas in drought induced CAM, core clock genes either change phase or amplitude, with novel CAM and stress specific cis-elements being responsible for rewired co-expression networks (Wai *et al*., 2019; Chen *et al*., 2020).

Plant species inhabiting deserts have been exposed to the so-called stressful environmental conditions for thousands of years. Many of them have developed morphological and physiological adaptations that enhance their photosynthetic performance and survival under extreme arid conditions (Gibson AC., 1998, Wang *et al*., 2019), including the evolution of CAM photosynthesis, CAM-C3 intermediate or C4 mechanisms (Heyduk *et al*., 2019, Wang *et al*., 2019, Folk *et al*., 2020). The Atacama Desert is one of the driest places on earth, with precipitations only occurring sporadically. When, every few years, thresholds on precipitation abundance and frequency are meet, the phenomenon known as the “blooming desert” emerge (Vidiella *et al*., 1999; Chávez *et al*., 2019; Araya *et al*., 2020; Holtum *et al*., 2021). *Cistanthe longiscapa* (Bernaud) Carolin ex Hershkovitz (Montiaceae) is an endemic annual plant and one of the most widespread and abundant plants within the blooming Atacama Desert events (Holtum *et al*., 2021). Previous reports based on leaf carbon isotope ratios suggest that some Chilean members of *Cistanthe* are in the C3-CAM intermediate spectrum (Arroyo *et al*., 1990; Palma and Mooney, 1998), with *C. longiscapa* representing a weak constitutive CAM (Holtum *et al*., 2021). Studies in other genus of the family, such as *Calandrinia* in Australia, have shown that many of these annual succulent species can be CAM facultative (Winter and Holtum, 2011; Holtum *et al*., 2017). In this work, we aim to explore the changes in gene expression associated to the C3-CAM spectrum in *C. longiscapa*, proposing that the intensity of CAM will vary concomitant with the intensity of aridity to which plants are exposed in the Atacama Desert. To this end, we studied the field variation on ecophysiological traits and CAM photosynthesis in *C. longiscapa*, assembled a *de novo* transcriptome followed by an RNA-seq comparative study from samples taken at dawn and dusk from three sampling sites, to track the changes in CAM/C3 switches under field conditions. The strength of this study lies in that it is the first transcriptomic study of an Atacama Desert blooming plant focused to unveil the molecular basis of the C3-CAM photosynthesis under natural environment conditions.

## Materials and Methods

### Sampling and plant traits measurement

Cistanthe longiscapa has a rosette with basal succulent leaves and inflorescence branches with showy terminal purple flowers, despite the strong impact of tourism during the blooming desert, this species is currently not threatened (Squeo *et al*., 2008). The study was conducted at the Atacama Desert, with all the samples being collected in July 2015, at the beginning of the flowering season. The year 2015 was a particular one in terms of precipitation timing, abundance, intensity (Wilcox. *et al*., 2016), and flowering abundance. Three sampling locations were selected between the localities of Copiapó and Vallenar, where populations of *Cistanthe longiscapa* (Bernéoud) Carolin ex Hershkovitz grows in extensive prairies when the flowering desert occur: Site 1 (S1) was set at 27°38’02.4”S 70°27’46.8”W, S2 at 27°57’57.6”S 70°33’25.2”W and S3 at 28°25’12.0”S 70°43’12.0”W. S1 and S2 site soils are calcisols, whereas site S3 display regosols (Supplementary Figure 1, Harmonized World Soil Database (version 1.2), Fischer *et al*., 2008). In each sampling site, fully expanded mature leaves from 10 healthy flowering plants were sampled at the evening (7-8 PM, henceforth “dusk”) and at the next morning (7-8 AM, henceforth “dawn”) on the same individuals, to determine nocturnal tissue acidification and prepare RNA extractions to cDNA library construction. Once sampled, all samples were immediately frozen in liquid nitrogen and transported to the laboratory in a Taylor-Wharton CXR500 dry shipper. Also, at dawn, leaves from the same plants were sampled to determine leaf mass per area (LMA), saturated water content (SWC), carbon isotope composition (δ^13^C (‰)), the percentage of Carbon and Nitrogen (%C, %N) and photosynthetic pigments content (chlorophyll A, chlorophyll B, carotenoids).

For LMA and SWC determination (Ogburn and Edwards, 2012), 5 leaves were freshly weighed, scanned for area determination, saturated with distilled water until constant weight, and oven dried at 75°C for 48-72 hours until reaching a constant weight. LMA was calculated as: Dry weight (g) / area (m^2^); and SWC as: [(Saturated weight (g) – dry weight (g)) / dry weight (g)]. For carbon isotope ratios and carbon and nitrogen composition, five fully expanded mature leaves per plant were dried, pooled, grinded, and analyzed at the Laboratory of Biogeochemistry and Applied Stable Isotopes (LABASI, PUC, Santiago, Chile) with an isotope ratio mass spectrometer and calculated against the Pee Dee belemnite (PDB) standard following Farquhar *et al*. (1989) equation. For chlorophyll and carotenoids content, leaves were instantly frozen in liquid nitrogen in the field and kept at - 80° C until the pigments were extracted with DMSO following the protocol and equations described by Wellburn (1994).

To estimate the degree of CAM photosynthesis, titratable acidity was measured with NAOH 0.01N (Keeley and Keeley, 1989) in leaves collected at dawn and dusk (10 plants per site). The accumulation was estimated as the difference between dawn and dusk and reported as nocturnal leaf acid accumulation. Significant differences for all measured traits were estimated using T-Test (for paired comparisons) or one-way ANOVA for each trait displayed. Principal component analysis (PCA) was used to evaluate correlations among traits. All the statistics were carried out in R (R Core Team, 2020).

### RNA isolation and cDNA library construction

Total RNA was extracted using Spectrum Plant Total RNA Kit (Sigma Aldrich, USA) from all the samples. We prepare 12 cDNA libraries (3 individuals □ 2 sampling times (dawn and dusk) □ 2 localities) using the TruSeq RNA-seq library prep kit from Illumina (Illumina, Inc., CA, USA) according to manufacturer’s instructions. cDNA libraries (Table S1) were sequenced in two lanes (paired-end 150 bp set-up) using HiSeq2500 sequencer (Macrogen Inc., Korea). Sequence data were deposited in the NCBI Short Read Archive in SUB10469409.

### *De novo* transcriptome assembly

Before the transcriptome assembly, FastQC was used to check reads raw quality. Raw reads were trimmed with the Trim Galore Cutadapt (Martin and Wang, 2011) wrapper using the paired and -q 25 option to conserve high quality reads quality. Overall quality was checked before assembly using MultiQC (Ewels *et al*., 2016). De novo assembly of high-quality contigs for *C. longiscapa* were performed using the Trinity package v2.5.0 (Haas *et al*., 2013) following the authors recommendations and with “min_contig_length 400” as additional parameter to avoid very short contigs. The assembled *de novo* transcriptome was optimized with the TransRate package (Smith-Unna *et al*., 2016) using the authors’ parameters recommendations. We look deeper into the transcriptome for ultra-conserved proteins using BUSCO (Simão *et al*., 2015) and the plant database Embryophyta odb09.

### *Cistanthe longiscapa* gene functional annotation

Gene functional annotation was performed using Mercator version 4 (Lohse *et al*., 2014) and EggNOG version 4.5.1 (Huerta-Cepas *et al*., 2016) to determine the best homologue from the model plant *Arabidopsis thaliana*. First, the predicted transcripts were converted into proteins using the Transcript decoder 2 tool from Cyverse (Joyce *et al*., 2017), using the universal genetic code and minimum protein length of 100 amino acids options. Transcript predicted proteins were next annotated using Mercator, using the following setting: TAIR release 10 database was used as reference, with a blast cut-off of 80 (default). The Mercator *A. thaliana* gene hits were annotated using the Thalemine (Krishnakumar *et al*., 2017) annotation. The same dataset was also assessed using EggNOG, using the following setting: DIAMOND for Mapping mode, Taxonomic Scope was adjusted automatically, Orthologs search was run using the “Restrict to one-to-one” mode, and Gene Ontology Evidence was run using the “Use experimental terms only” mode.

### Principal component and differential gene expression analysis

To maximize differences, only RNAseq from individuals from the extreme populations (S1 and S3) were used for differential expression analysis. Principal Component Analysis (PCA) was carried out using the R package FactoMineR (Lê *et al*., 2008). Differential expression analysis was performed for libraries from S1 and S3 between conditions (dawn and dusk), using the DESeq2 package (Love *et al*., 2014). Counts data was pre-processed to keep genes with at least 1.0 counts per million (cpm) in at least 3 samples. To maximize the number of differentially expressed genes detected, while controlling the false discovery rate (FDR), the Bioconductor package Independent Hypothesis Weighting (IHW, Ignatiadis *et al*., 2016) was used to determine the adjusted *p-value* on DESeq2. Volcano plots were generated using the ggplot2 R package whereas heatmaps were generated using the pheatmap R package.

### Gene ontology and pathways enrichment analysis

Gene ontology (GO) annotation from the best *A. thaliana* homologues was retrieved from the PlantGSEA (Yi *et al*., 2013) “A. thaliana GO gene sets”, and transferred to the genes from *C. longiscapa*. Next, this mapping was used to generate a Gene Matrix Transposed (GMT) file, where each row maps a GO term to the genes from *C. longiscapa* which have this annotation. This GMT file was used to assess GO enrichment using the g:Profiler web server (Raudvere *et al*., 2019). In order to better characterize the g:Profiler results, the enriched GO terms redundancy was removed using REVIGO (Supek *et al*., 2011), with the following parameters: Allowed similarity: Small (0.5); GO categories associated to: P-values; GO term sizes database: *A. thaliana;* semantic similarity measure to use: SimRel. Next, all the GO terms displayed by REVIGO were further summarized using the mclust method available at the simplifyEnrichment R/Bioconductor package version 1.2.0.

Pathway’s enrichment analysis was also performed using the g:Profiler web server, based on the in house generated ClongiscapaCyc PTOOLS v25.0 pathway-genome database GMT file (PGDB, Karp *et al*., 2021), build using as input the annotation results generated by the e2p2v4 enzymes annotation tool (Schläpfer *et al*., 2017).

### Gene regulatory networks analysis

DESeq filtered and normalized counts transcripts (Supplementary File 1) were used for inferring and analyzing gene regulatory networks (GRNs) related to the S1 and S3 site plants, using the R package DIANE (Cassan *et al*., 2021). Known transcriptional regulators related to CAM photosynthesis were retrieved from the literature (Supplementary Table S1, Brilhaus *et al*., 2016, Amin *et al*., 2019, Maleckova *et al*., 2019, De La Harpe *et al*., 2020, Moseley *et al*., 2021). After inference of the networks and empirical p-values assessment of the regulator-gene pair weights, edges above an FDR of 0.05 were kept to generate the final networks (Cassan *et al*., 2021).

### qPCR validation

We validated, by qPCR, some of the transcripts that were differentially expressed among libraries and were related to “CAM photosynthesis”, “Circadian Rhythm” and “Abiotic Stress” in the same samples used to perform the RNA-Seq libraries. One microgram of total RNA was treated with 1μL of DNAse I, amplification grade (Thermo Fisher Scientific) according to the manufacturer’s instructions. This DNA-free RNA was used as a template for first-strand cDNA synthesis with an oligo(dT) primer and SuperScript II (Thermo Fisher Scientific), according to the manufacturer’s instructions. The primers described in Supplementary Table S2 were used to amplify PCR products from single-stranded cDNA. qPCR was performed using the Fast EvaGreen qPCR Master Mix kit (Biotum, CA, USA). Reactions contained 1 μL of 1:10 diluted cDNA in a total volume of 10 μL. The quantification and normalization procedures were done using the following equation (Vandesompele *et al*., 2002; Hellemans *et al*., 2007):

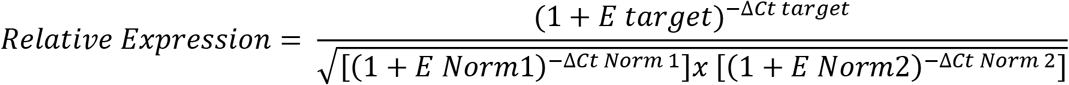

where E corresponds to the efficiency of amplification of target and reference genes, Ct is the threshold cycle, and Norm1 and Norm2 refer to the references or normalizer genes. We selected *ClClathrin* and *ClGAPDH* as normalizer genes.

## Results

### *Cistanthe longiscapa* sampling on the Atacama’s Desert

*C. longiscapa* (Figure 1, upper and mid panels) extends its distribution between 25° to 31°S, from coastal habitats up to 3,800 m of elevation in sandy soils (Hershkovitz, 1991), where mean annual precipitation modeled after Bioclim 2.0 (Fick and Hijmans, 2017) is less than 40 mm/yr (Figure 1, lower panel). This species is widely distributed in valleys and coastal plains during the blooms after rare winter rainfall in the Atacama Desert (Araya *et al*., 2020; Holtum *et al*., 2021). The flowering period usually extends between September and December (late winter to early summer). Plants were collected from three sites along the Atacama Desert environmental arid gradient (López *et al*., 2016), displaying differences in precipitation values (Figure 1, lower panel) and soil type (Supplementary Figure 1).

**Figure 1.**
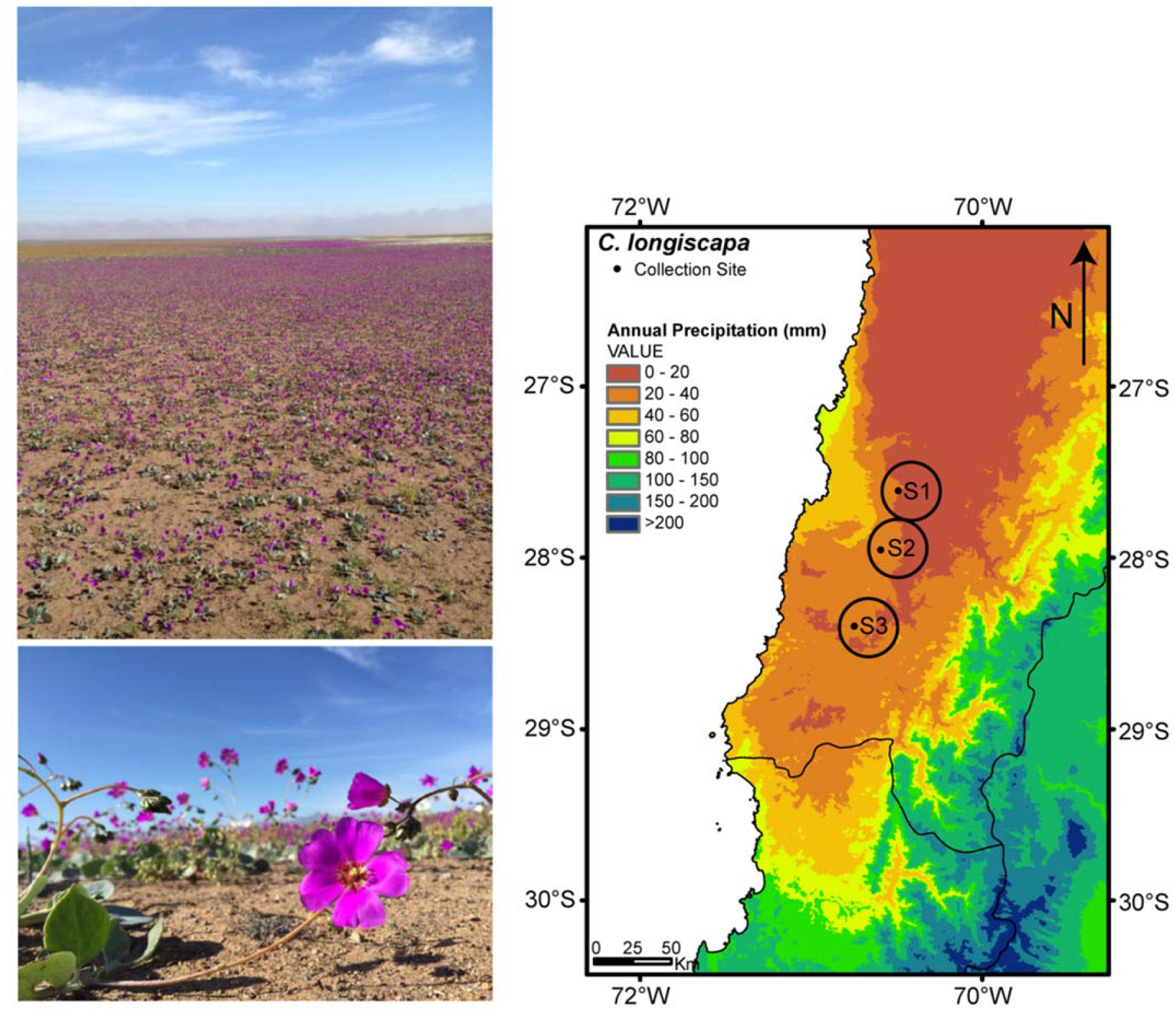
*Cistanthe longiscapa* species and collection sites description. The species model *Cistanthe longiscapa* can be seem as plants distributed as mantles of purple flowers in contrast with the arid Atacama Desert soil (upper and mid panels). A closer look allows the identification of a rosette with basal succulent leaves and inflorescence branches with terminal purple flowers. The geographic location of the study sites (S1, S2 and S3, lower panel) mapped against the average annual precipitation was obtained using Bioclim 2.0.

### Ecophysiological leaf traits points to different photosynthetic processes between S1 and S3 plants

To determine whether plants from the different sites displayed different ecophysiological performance, seven parameters associated with CAM and photosynthesis were analyzed (Figure 2A, Supplementary Figure 2). To this end, nocturnal leaf acid accumulation, isotopic ratio (δ13C), photosynthetic pigments (Chla, Chlb, Carotenoids), Carbon to Nitrogen ratio (C/N), leaf mass per area (LMA) and succulence (SWC) were assessed in leaves from ten plants from each site. Regarding the nocturnal acid accumulation, the highest values were reached in the northern site (S1, Δ Acidity = 251.56 ± 43.79 meqH+/g) and the lowest in the southern site (S3, Δ Acidity = 108.18 ± 41.8 meqH+/g), covering a spectrum of CAM photosynthesis intensity from strong to weak towards the south. The isotopic ratio (δ13C) measured in leaves of *C. longiscapa* also supports this spectrum, ranging from −15.42 to −24.15 ‰, values indicative of strong to weak CAM or C3-CAM intermediate (Messerschmid *et al*., 2021). For instance, the average in plants from S1 support a strong CAM (δ13C = −17.5‰ ± 1.2) in this northern sampling site, whereas the mean value in plants from S2 and S3 are more in agreement with weak CAM or C3-CAM photosynthesis (S2 δ13C = −22.2‰ ± 1.1; S3 δ13C = −19.5‰ ± 1.1). Photosynthetic pigments (Chla/Chlb = 2.13 ± 0.35) and Total Chlorophyll/Carotenoids (Ctot/Car = 3.41 ± 0.51) showed their lowest values in plants from S1, increasing towards S3, an opposite pattern compared to the nocturnal leaf acidification. Plants from the S1 site had the lowest leaf mass per area (131.35 ± 28.76), whereas the highest value for this parameter was in plants from S3 (170.84 ± 23.94). Regarding the Carbon to Nitrogen ratio, the highest value was found in plants from S1 (17.88 ± 2.13), whereas the lowest level was found in plants from S3 (12.55 ± 2.63). Finally, succulence (SWC) was higher in leaves from plants from the S1 compared to S2 and S3 sites.

**Figure 2.**
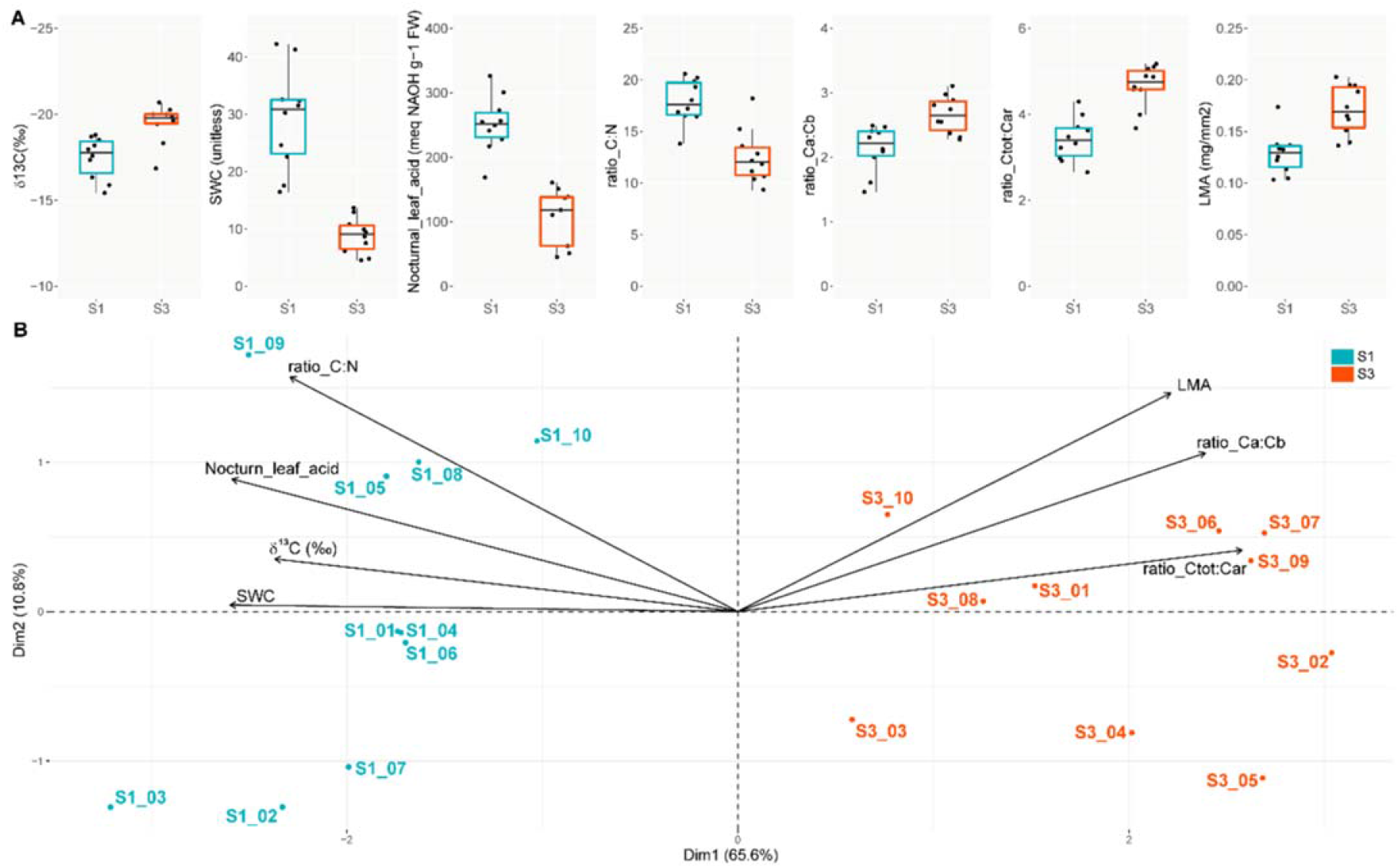
Relation among ecophysiological variables measured in individuals from 2 different sites across the Atacama Desert. Differences among sites S1 and S3 plants, in terms of nocturnal acid accumulation (Nocturnal_leaf_acid), isotopic carbon ratio (δ^13^C), leaf mass per area (LMA), succulence (SWC), total Chlorophyll/Carotenoids (ratio_Ctot:Car), Carbon to Nitrogen ratio (ratio_C:N) and photosynthetic pigments ratio (ratio_Chla:Chlb), were assessed (**A**). Principal component analysis using the data from these 7 traits allowed segregating and correlating the sampling sites with the different ecophysiological variables (**B**). TTtest, was used to assess the differences between sites S1 and S3 plants. All comparisons p-values were smaller than 0.005. N = 10.

To evaluate if the ecophysiological variations observed among the plants from the different sites replicate the geographic samples distribution, we performed a PCA analysis (Figure 2B). We observed that some variables strongly explain the segregation among geographic sampling sites. For example, the three best explanatory traits for CAM, δ13C, nocturnal leaf acidity accumulation and succulence (SWC) were correlated with Dim 1 (which accounted for 48.4% of the variance) and S1 site, segregating those samples from the other two sites. Leaf mass per area (LMA) and photosynthetic pigments (ratio Chla/Chlb and Ctot/Car) were correlated with Dim 1 in the opposite direction, and with S3 samples, explaining their segregation towards the other extreme of the Dim1 axis. These results suggest that *C. longiscapa* plants located in S1 and S3 sites performed different photosynthetic mechanisms.

### *De novo* assembly of the *Cistanthe longiscapa* transcriptome and differential gene expression uncover different stress responses in plants from sites S1 and S3

To get insights of the transcriptome repertoire of *C. longiscapa*, a de novo assembly was carried out using RNA-seq reads from 12 libraries obtained from sites S1 and S3, collected at dusk and dawn (Table 1), with a contig N50 of 1,128 bp. A total of 88,770 transcripts were predicted, which could be translated into 43,241 non-redundant proteins. The comparison against a set of highly conserved genes using BUSCO resulted in 61% of completed and 14% of fragmented genes, while 25% of the genes were missing. Close to 90% (88.8%) of the predicted proteins were associated to a homologous protein when using in parallel Mercator (Lohse *et al*., 2014) and EggNOG (Huerta-Cepas *et al*., 2016) gene annotation tools (Supplementary Table S3). When considering only the *A. thaliana* best homologues, annotated by Mercator, 32,886 transcripts (76.1%) were covered, with a one-to-one relationship for 15.5% of these transcripts.

**Table 1:** Assembly and annotation statistics for *Cistanthe longiscapa*.

| Parameters | Values |
| --- | --- |
| Trinity genes | 88770 |
| GC (%) | 42 |
| Median contig (bp) | 734 |
| Average contig (bp) | 990 |
| Contig N10 (bp) | 2926 |
| Contig N20 (bp) | 2191 |
| Contig N30 (bp) | 1731 |
| Contig N40 (bp) | 1397 |
| Contig N50 (bp) | 1128 |
| Minimum length (bp) | 401 |
| Maximum length (bp) | 402673 |
| Annotated (*) | 50280 (56.64%) |
| BUSCO (**) | 61 (C); 14 (F); 25 (M) |

(\*) Aligned under non-redundant database (nr)
(\*\*) Percentage of a total of 1440 genes, (C) Complete; (F) Fragmented; (M) Missing

Principal component analysis of the RNAseq sample libraries clearly shows that the first dimension discriminates by site of collection whereas the second discriminate by time of sample collection (Figure 3A). Together, both dimensions accounted for 42.4% of the variability in the transcriptome. In terms of data quantification, a total of 69,614 transcripts of the *de novo C. longiscapa* transcriptome had expression values above 0. After the removal of lowly expressed and low reproducibility genes, the gene set to be assessed accounted for 35.3% of the original transcriptome (Supplementary Table S4, Supplementary File 1). Genes with differential gene expression added up to 3,991 when comparing dawn versus dusk at S1, and 2,564 at S3 (Figure 3B and C), with both sites predominantly displaying higher gene expression levels at dawn (Figure 3D). At S1, the dawn/dusk ratio was 2.3 (2771/1220 transcripts), whereas at S3 the ratio was 1.4 (1491/1073 transcripts). Regarding the genes that were more expressed at dawn in both sites, close to 24% were shared (822 transcripts), whereas genes that were more expressed at dusk in both sites were close to 18% (343 transcripts, Figure 3D).

**Figure 3.**
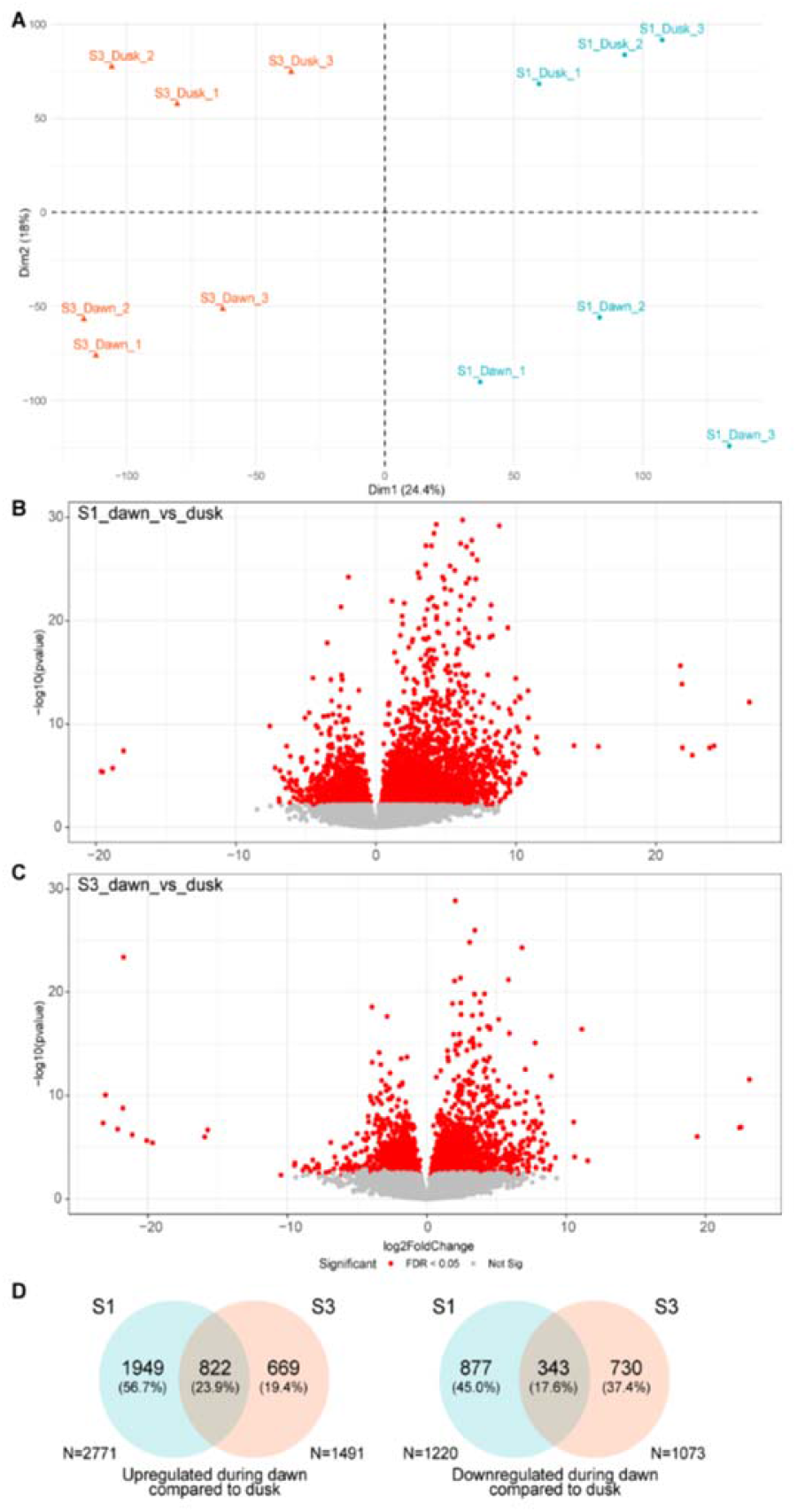
Relation *Cistanthe longiscapa* field data transcriptome analysis. Data from the *de novo* transcriptome from S1 and S3 plants, collected at dawn and dusk, was characterized by means of principal component analysis (**A**). The first dimension (PC1) split the data from sites S1 and S3, whereas PC2 split the data into dawn and dusk samples. Replicates, as expected, clustered together. The number and magnitude of transcripts significantly up or down-regulated at dawn compared to dusk are depicted as volcano-plots (**B**). At both S1 and S3 sites, most transcripts were upregulated at dawn, with the Site S1 displaying the higher number of transcripts differentially expressed. The maximum rates of change in both sites are similar, as can be seen by the units of the log2FoldChange X axis. In terms of similarities, sites S1 and S3 displayed 23.9% of the genes with similar up-regulation and 17.6% with similar downregulation at dawn (**C**).

Functional enrichment analysis of this data was performed using the Gene Ontology (GO) annotation (Figure 4A; Supplementary Table S5) and MetaCyc pathways (Figure 4B). GO-based analysis allowed the identification of 64 processes enriched in some of the four conditions assessed. By using the corresponding semantic similarity matrix of these 64 GO terms, we further grouped them in 8 clusters (Supplementary Figure 3). Clusters 1, 2 and 6 were similarly represented in both sites, whereas cluster 3 displayed an opposite pattern for each site, being related to circadian processes. Cluster 4, related to cell death, was only associated with S1 plants at dawn. Cluster 5 was more diverse and included GO processes related to photosynthesis associated with genes with higher expression at dawn at the site S3. Clusters 7 and 8 were related to abiotic stress, being preferentially associated to the transcripts with higher expression at dawn at the site S1.

**Figure 4.**
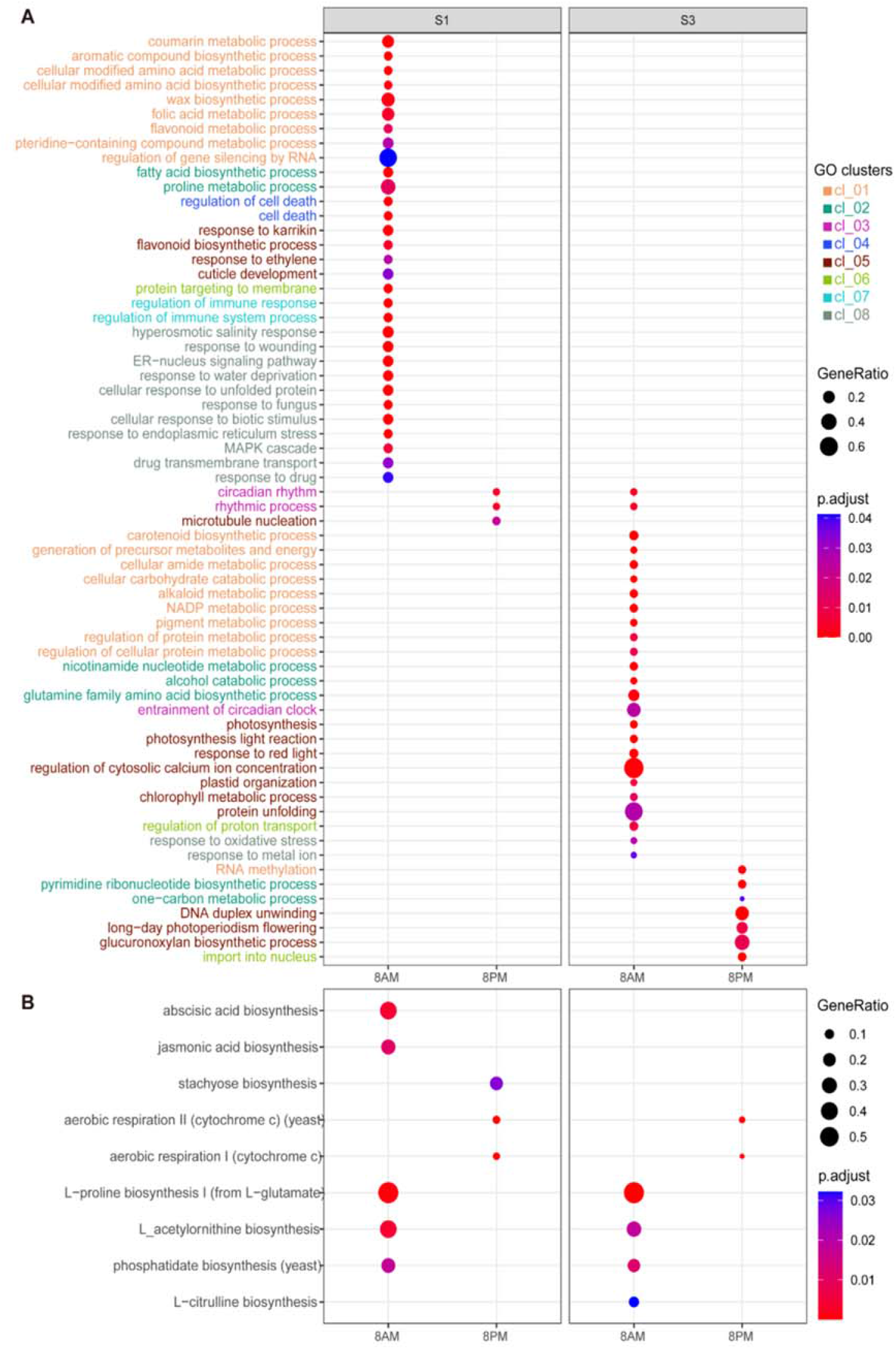
Functional enrichment analysis of genes upregulated at dawn or dusk in sites S1 and S3. The ratio and significance of GO (**A**) and MetaCyc pathways (**B**) of genes enriched at sites S1 and S3 at 8 AM (up-regulated at dawn) and 8PM (up-regulated at dusk) are shown as dotplots. GO biological process (BP) terms were further grouped in 8 clusters according to their semantic similarity (Methods).

In terms of pathways, abscisic acid biosynthesis displayed many genes with higher expression at S1 at dawn (Figure 5A), with genes involved in the biosynthesis of intermediate compounds and ABA transport also upregulated. Another set of pathways with a different expression profile between S1 and S3 were highlighted in Figure 5B. These pathways are key for regulating glutamine and glutamate accumulation and are also responsible for the biosynthesis of γ-aminobutyrate (GABA) which, as well as ABA, prompt plants to close stomata, a process that would be triggered at dawn in S1 site plants.

**Figure 5.**
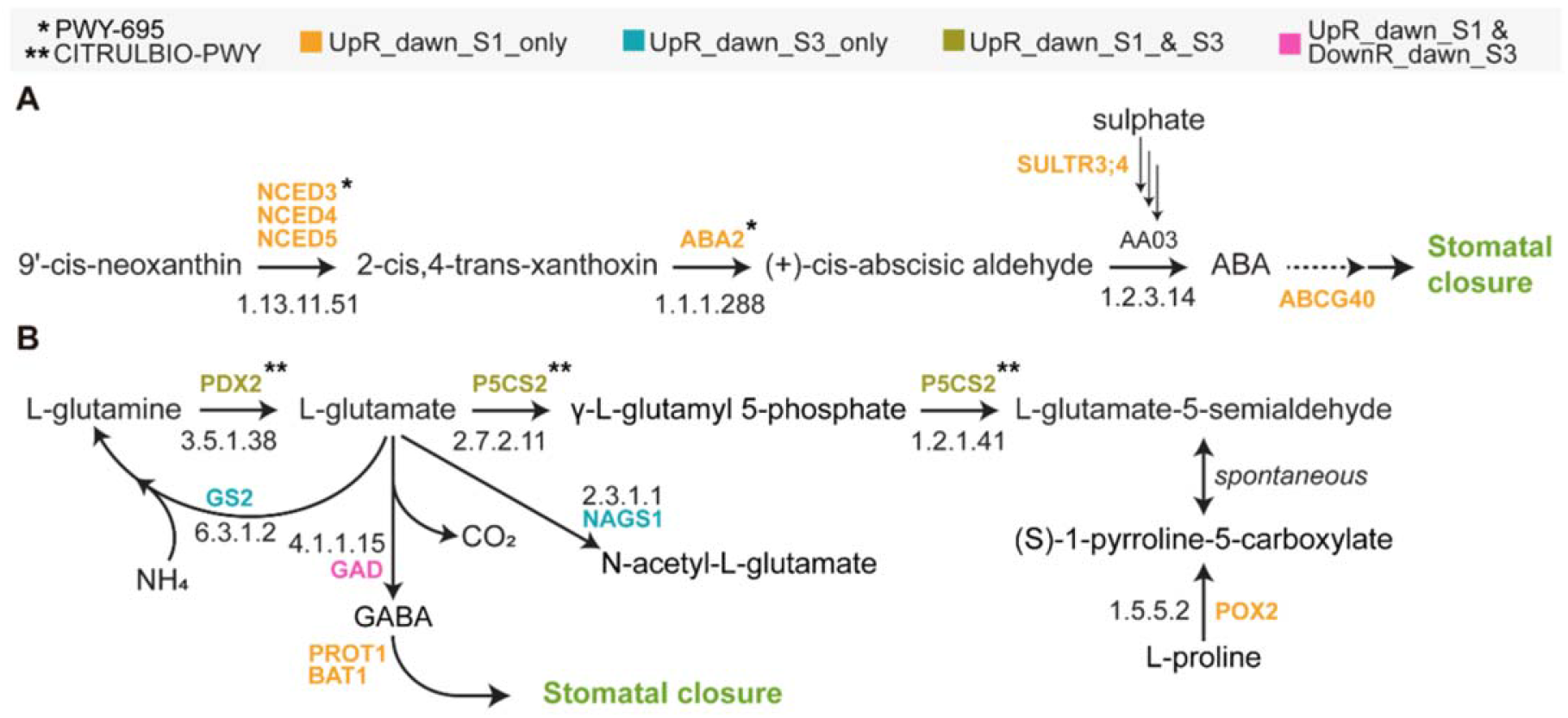
Pathways related to stomatal closure. The ABA biosynthesis pathway (**A**, PWY-695) included several genes up-regulated at dawn at the site S1 (stressed in bold-orange). We also pointed out genes that displayed the same pattern and that could be involved in ABA transport or in the biosynthesis important precursors for the enzymes of the pathway activity. The other pathways assessed were the L-citrulline biosynthesis (CITRULBIO-PWY), the acetylornithine biosynthesis (PWY-6922) and the proline biosynthesis I (PROSYN-PWY) pathways (**B**). Genes upregulated at dawn in both sites, or in S1 or in S3 are highlighted with different colors. Interestingly, the result of this regulation of these pathways would be the stomatal closure during the day in S1 site plants. ABA - abscisic acid; GABA - γ-aminobutyrate.

### Gene regulatory network analysis identifies the gene network of S1 plants up regulated at dawn as the most complex

An analysis of 50 genes related to the control of the circadian clock (Supplementary Tables S6 and S7, Moseley *et al*., 2021) showed that 28% of these genes displayed a higher expression at dawn, led by LHY1, which peaks in the morning, and the other 20% displayed a higher expression at dusk and were led by TOC1, which peaks in the evening (Schiller and Bräutigam, 2021). S3 plants displayed 20% of the 50 assessed circadian genes with an exclusive upregulation at dawn, as opposed to S1 plants, which displayed other 20% of the circadian clock genes with an exclusive upregulation at dusk.

Several of the genes involved in the control of the circadian clock have been recognized as key CAM transcriptional regulators (Brilhaus *et al*., 2016, Amin *et al*., 2019, Maleckova *et al*., 2019, De La Harpe *et al*., 2020, Moseley *et al*., 2021). Therefore, we assessed the differences between S1 and S3 plants in terms of gene regulatory networks. The S1 GRN at dawn displayed more regulatory connections than any other GRN, with a link density of 1.142 compared to 0.980 from the S3 GRN at dawn, the network with the second larger number of regulatory connections. Among the hubs found, the nuclear-encoded sigma factor 5 (SIGE), the *A. thaliana* homologue of AT2G44730, and the heat shock transcription factor A2 (HSFA2) were the top 3 (Table 2).

**Table 2.**
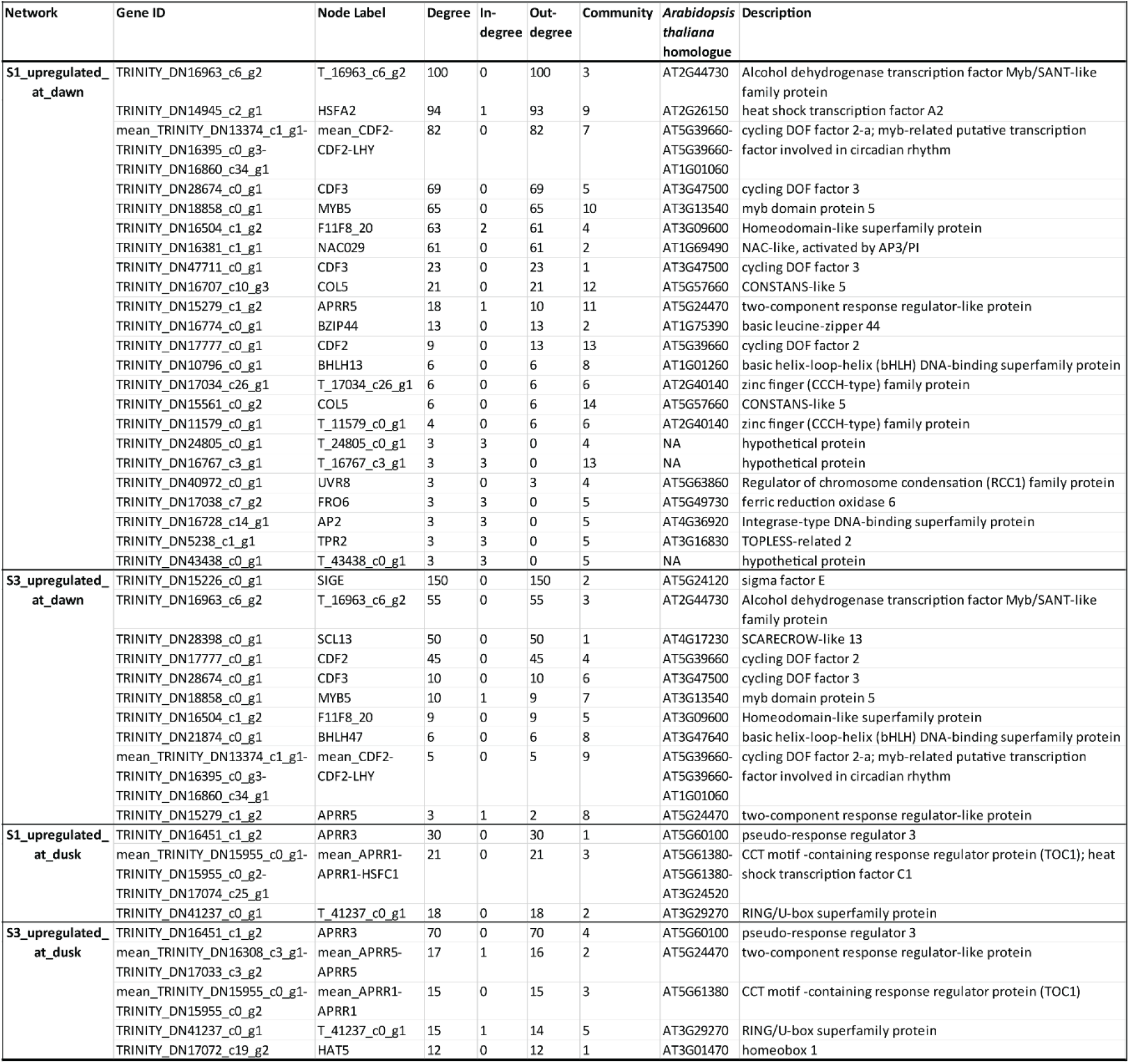
Gene Regulatory Network hubs associated to S1 and S3 plants differentially expressed genes.

Community assessment based on GO enrichment analysis and normalized expression profiles (Supplementary Figure 4) also allowed to discriminate similarities and differences among the GRNs. For instance, the community number 3 in both S1 and S3 plants, at dusk, were similar in size, biological process GO enrichment and main hub genes. The community number 2 of the genes upregulated at dawn in S3 plants was, in turn, related to photosynthesis and displayed an expression change only in S3 plants. In addition, its main hub was the chloroplast-localized sigma factor SIGE, an essential factor in the nuclear control of chloroplast function and its response to environmental stress (Mellenthin *et al*., 2014, Zhang *et al*., 2020).

### Validation of Differential Expression of Genes using qPCR

Based on the ecophysiological response and the enrichment on several GO terms associated to photosynthesis on plants from the site S3 at dawn, we evaluated the transcript level of genes associated to CAM metabolism such as PEPCK, PPDK, NADP-ME4 and RUBISCO activase (Figure 7). Among them, the transcript level of NADP-ME4 accumulates at dawn on plants from both sites whereas the transcripts of PEPCK, PPDK and RUBISCO activase did not shown a particular pattern of expression between the sampled sites, although PPDK transcript accumulation was higher on S1 samples. Given the fact that GO terms associated to circadian rhythm displayed a contrasting temporal pattern between sites S1 and S3 (Figure 4), the transcript level of two genes associated the dark phase such as APRR1, Gigantea and one gene associated to the light phase like Late Elongated Hypocotyl (LHY) were evaluated. As expected, the transcript levels of APRR1 and Gigantea reached their maximum at dusk, whereas the transcript of LHY reached their maximum at dawn and the level of all transcripts were higher on samples from site S3 (Figure 7). Additionally, we evaluated the transcript levels of genes associated to ABA biosynthesis, NCED3 and ABA2, considering the GO terms and MetaCyc pathways enrichment in response to water and plant hormone signaling transduction (Figure 5). As shown on Figure 7, the transcript level of NCED3 was higher on samples from S1 at dawn whereas the transcript of ABA2 exhibited similar levels at dawn and dusk at each sampled site, but with a slightly higher amount on S1 samples.

## Discussion

The Atacama Desert is a strongly arid environment, with a trend of aridity intensification towards the northern hyperarid core, particularly between the latitudes 30° and 25°, which enclose our sample sites (Figure 1, López *et al*., 2016). The stochastic and rare rain-driven Atacama Desert blooming episodes can generate spatially isolated plant communities that remain separated during the intervening dry periods (Holtum *et al*., 2021), providing a well-suited condition for functional traits intra-specific divergence according to the microenvironments the species are exposed to. Studying intra-specific variation can help understand evolutionary adaptations to environmental change and the plants stress response (May *et al*., 2017). In this study evaluated intra-specific functional traits from the annual plant *Cistanthe longiscapa*, a conspicuous member of the Family Montiaceae, that grows during the “blooming desert” events. Plants located at different desert landscapes (Figure 1 and Supplementary Figure 1) were ecophysiologically and genetically characterized, which allowed us to detect different levels of CAM photosynthesis in *C. longiscapa* plants from different sites. Focusing on the transitional phases II (dawn) and IV (dusk) of the four-phase diurnal CAM cycle (Osmond CB, 1978), we were able to capture transcriptional information from key processes for the plant response to the environment, such as plant photorespiration, ABA and GABA biosynthesis and circadian regulation, which could help understanding the molecular basis for the metabolic flexibility shown between *C. longiscapa* plants inhabiting environments with differences in aridity (Winter K, 2019).

CAM plants can be characterized by displaying a nocturnal acidification, a feature not found in C3 plants (Winter and Smith, 2022). In this study we were able to detect different levels of nocturnal leaf acidification in *C. longiscapa* plants collected from different sites and to associate these differences with the development of an abiotic stress response. These variations in acidity correlate with differences in δ13C contents, especially between the most northern (S1) and southern (S3) collection sites (Figure 2), pointing to a constitutive CAM in this species (Holtum *et al*., 2021), but with an environmentally sensitive CAM modulation (Winter K., 2019, Schweiger *et al*., 2021). In addition to the differences of δ13C and nocturnal titratable acid accumulation between S1 and S3 sites, our ecophysiological results disclose higher chlorophyll a/b and chlorophyll/carotenoids ratios in S3 plants, which could represent positive adaptive values of plants performing photosynthesis under high light stress conditions (Gori *et al*., 2021). Differential expression of genes related to photosynthesis and photorespiration at S3 also support the C3 diurnal unfolding in S3 plants (Figure 4A). These results are remarkable because they show that *C. longiscapa* is a species capable to live under extreme conditions of water scarcity by using a resource conservative strategy such as CAM, that could be switched to a CAM-C3 resource expenditure strategy in response to almost imperceptible changes in arid conditions that would provide a more amenable environment (Pereira and Cushman, 2019).

Other leaf functional trait analyzed was “Leaf mass per area” (LMA), that is the ratio between leaf dry mass and leaf area. This trait would account for carbon and nutrients that are invested in a certain area of light-intercepting foliage, reflecting the leaf-level cost of light interception (Poorter *et al*., 2009), that is, high structural investment, lower mesophyll conductance. In a strictly C3 plant we would have expected lower LMA values associated with higher C3 photosynthesis performance but our results showed that plants from S1 have the lowest LMA values, where carbon isotopic and leaf acidity values were indicatives of more CAM and lesser C3 photosynthesis performance. This counterintuitive pattern can be explained when we broke down LMA into leaf volume to area ratio (LVA, mL m-2) and leaf density (LD, g mL-2) (De la Riva *et al*., 2016). Because of the presence of large volumes of water-storage cells in succulents, LVA is over 10 times higher in succulent than in non-succulent species, driven the variations in LMA in this kind of plants (Nielsen *et al*., 1997). The negative correlation between LMA and succulence in our dataset (Figure 2) corroborate this rationale, indicating that the observed variaton trend in LMA within these plants is due to variation in water storage rather than structural investment. In small annual plants, such as *C. longiscapa*, succulent leaves would represent single-use water stores designed to extend the growing season into the portion of the year where resources such as water become scarce (Males J., 2017). If the variation in LMA is due to LVA rather than LD, it is expected that Carbon content remains relatively constant among sites. In effect, C:N ratio remains constant in S1 and showed a sharply decreased in S3 (Supplementary Fig. S2).

CAM nocturnal carbon uptake is made possible by the inverse stomatal behavior compared to C3 plants, meaning stomatal opening at night and closure during all or part of the day, leading to reduced water loss and improved water-use efficiency. Stomatal aperture is regulated by diverse environmental signals, such as light and CO_2_, as well as by internal plant signals, such as abscisic acid (ABA) (Schiller and Bräutigam, 2021). Pathway-enrichment analysis indicates that S1 plants display an enrichment in ABA and jasmonic acid (JA) biosynthesis pathways (Figure 4B). ABA signaling can be induced by drought, followed by stomatal closing to prevent water loss (Schiller and Bräutigam, 2021).

Therefore, ABA synthesis and transport to target sites would be particularly required for plants performing CAM photosynthesis, such as S1 plants. Noteworthy, we also found that sulphate transport might be regulated (Figure 5A) to increment ABA levels, possibly by providing sulfur for cysteine biosynthesis, which can be used downstream by ABA3, a MoCo-sulfurylase required for activating AAO3, the last step in the ABA biosynthesis (Cao *et al*., 2014). Also regulating AAO3 would be one of the main hubs of the S1 plant genes upregulated at dawn, the NAC transcription factor 29 (NAC29, Figure 6). NAC29 is, in fact, a TF associated with cold and drought responses that would have shown a lineage-specific gene expansion in the *C. longiscapa* family, the Montiaceae (Wang *et al*., 2019).

**Figure 6.**
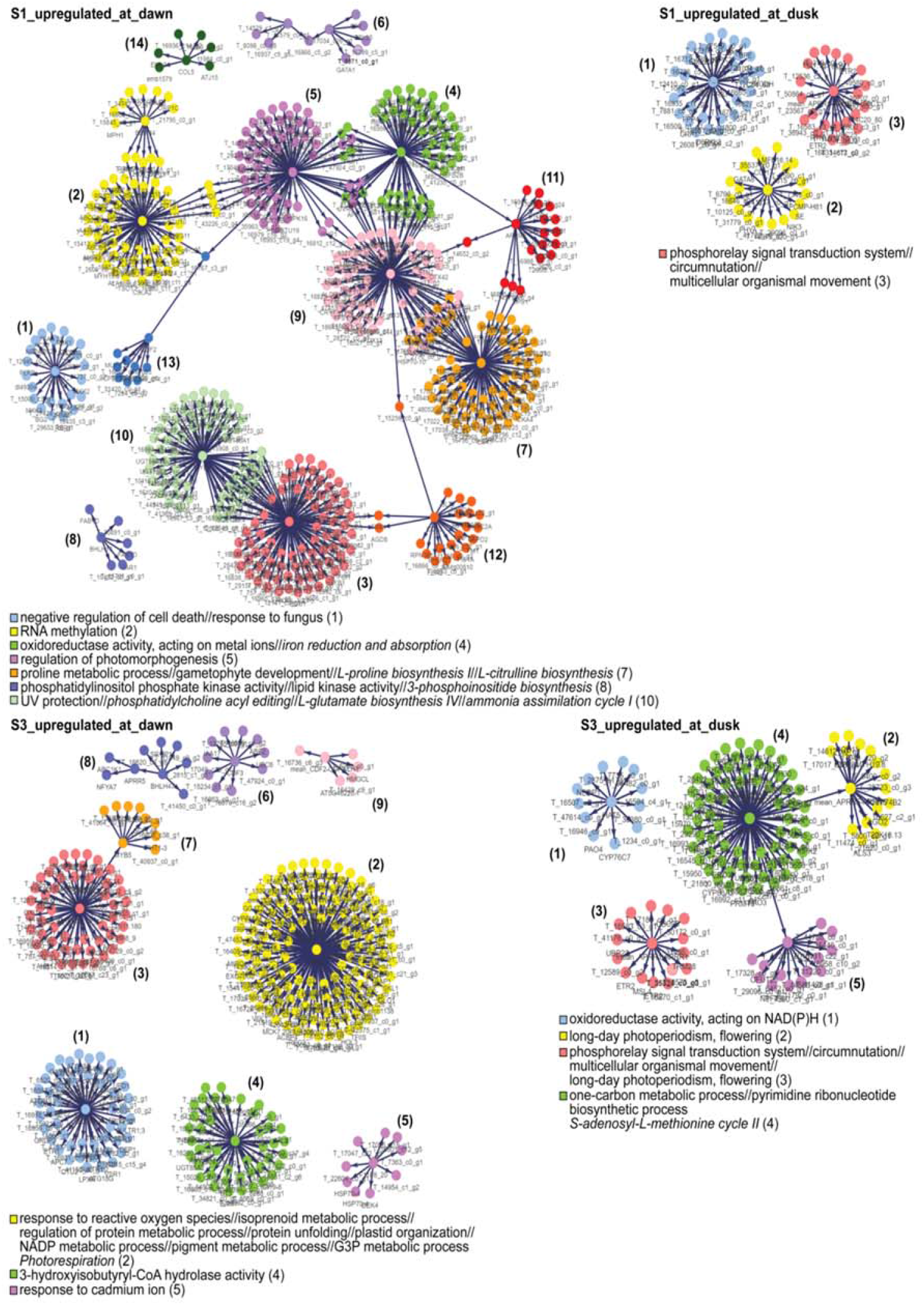
Gene regulatory networks associated to the expression profiles of S1 and S3 plants at dawn and dusk. GRNs were generated using the genes expression profile from S1 and S3 plants, at dawn and dusk. S1 and S3 networks at dawn were composed by many more communities (topological modules) than at dusk. The results of the communities GO enrichment analysis are shown below each network, with colors and numbers for each community used as identifiers. Enriched pathways are also indicated in italics. GO terms and pathways associated to the same community annotation are separated by a doble slash (//). Notice that the community colors are not related to the genes they are composed of, they just allow discriminating then.

**Figure 7.**
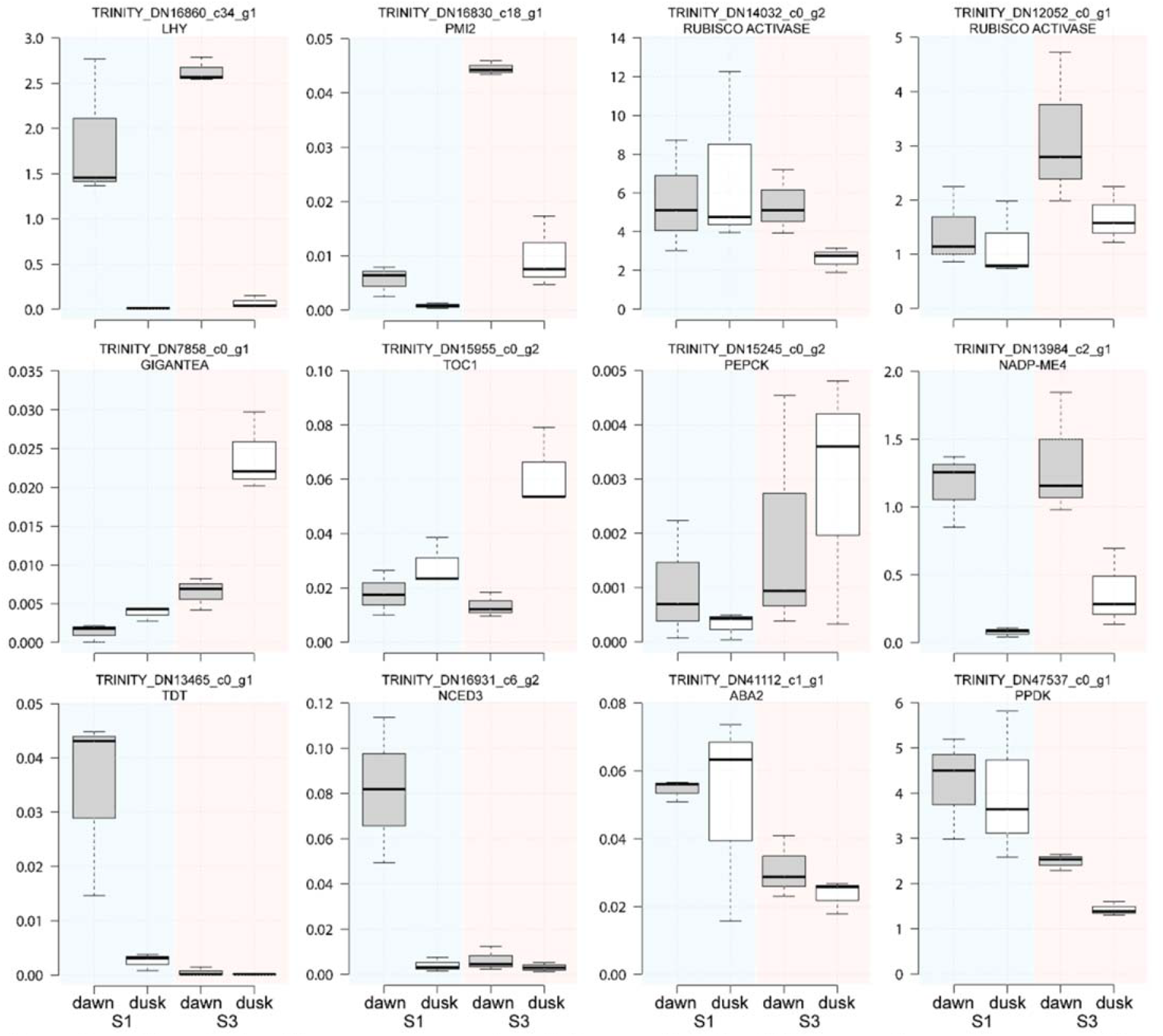
qPCR analysis of key genes expression at dawn and dusk. qPCR was performed for some of the transcripts that were related to “CAM photosynthesis”, “Circadian Rhythm” and “Abiotic Stress” in the same samples used to perform the RNA-Seq libraries. The primers used are described in **Supplementary Table S2**.

Many signaling pathways, besides those related to ABA, can be involved in regulating stomatal opening under drought stress conditions (Askari-Khorsgani *et al*., 2018). When assessing the differences between plants from the S1 and S3 sites related to pathways involved in the metabolism of key metabolites for the plant nitrogen homeostasis such as glutamine and glutamate (Figure 5B), it is possible to foresee how plants from the S3 site could use the enzyme glutamine synthetase 2 (GS2) at dawn to recycle ammonia derived from photorespiration (Ferreira *et al*., 2019). S1 plants, in turn, would be using the enzyme glutamate decarboxylase (GAD) to generate CO_2_ and GABA, a compound known to be involved in reducing stomatal opening, therefore improving water use efficiency and drought tolerance (Xu *et al*., 2021), as well as generating diurnal CO_2_ (Carillo P., 2018) under conditions where atmospheric CO2 would be limiting. In summary, the differential accumulation of the metabolites ABA and GABA could have a key role in the CAM diurnal stomatal closure of S1 *C. longiscapa* plants.

The inversed temporal CO_2_ fixation characteristic of the CAM photosynthesis is apparently correlated to changes in the plant circadian clock (Wai *et al*., 2019, Moseley *et al*., 2021, Schiller and Bräutigam, 2021). *Sedum album* plants performing C3 and CAM drought-induced photosynthesis displayed 12.9 and 18.6% of the assessed genes cycling in one condition or the other, with only 22% of the genes cycling in both conditions (Wai *et al*., 2019), highlighting the significative rewiring of the transcriptome in response to drought conditions. About 48% of the genes related to the control of the circadian clock assessed in this work displayed a conserved pattern in S1 and S3 plants. S3 plants displayed 20% of the assessed circadian genes with an exclusive upregulation at dawn, compared to S1 plants. Among those were PHYB and CRY2 which, together, regulate the chromatin degree of compaction under low light conditions (Martínez-Garcia and Moreno-Romero, 2020), enabling the transcription of the genes located in the exposed euchromatin regions.

A fine temporal and spatial regulation of metabolites and gases flow is required in CAM to avoid futile cycles, to favor the proper metabolites storage and to overcome the obstacles imposed by the succulent CAM plants leaf architecture. However, there are still several transporters that are unknown and are key for a CAM plant operation (Winter and Smith, 2022). Among the many transporters with a differential expression at dawn in S1 plants, we found several clearly related to the CAM metabolism: Dicarboxylate transporter (DiT1) and an aluminum-activated malate transporter (ALMT1) could be involved in the export of malic acid from the vacuole at dawn (Wai *et al*., 2019, Ceusters *et al*., 2021).

Succulent tissues might be advantageous for CAM plants due to the possibility to maximize the acid- and water-storage capacity of its cells (Lim *et al*., 2020, Winter and Smith, 2022). *C. longiscapa* plants from the S1 site, who would be performing CAM metabolism, displayed on average 3 times larger succulency compared to site S3 plants (Supplementary Figure 1). However, succulent leaves present tight packed water-rich cells, which represents an obstacle for CO_2_ diffusion due to the lower diffusion speed of this gas in water compared to air (Schiller and Bräutigam, 2021). Aquaporins have been identified as facilitators of CO_2_ diffusion across membranes (Gago *et al*., 2020), and therefore are good candidates for improving CO_2_ diffusion in CAM succulent leaves. S1 plants displayed a higher level at dawn of two PIP1 proteins homologous to PIP1A and B, which could help the diffusion of CO_2_ across the plant leaf tissues (Heckwolf *et al*., 2011).

## Conclusions

The understanding of how a species can adjust its metabolism to perform C3 or CAM photosynthesis as a response to changes in its environment is key for CAM engineering in C3 crops, since it provides the opportunity to trigger CAM photosynthesis in periods of time when the plants are more susceptible to drought, and then return to a less energetically expensive C3 mode. *C. longiscapa* is an annual “blooming desert” species that can perform such transition as established in the present study, based on ecophysiological and transcriptomic analysis of field samples.

*C. longiscapa* plants performing CAM photosynthesis would rely on the phytohormone ABA and the signaling molecule GABA to reduce stomatal opening, and in more succulent leaves to provide the proper leaf’s vacuolar storage capacity required for performing CAM (Töpfer et al., 2020). The temporal regulation of the processes that allow the switch between a weak into a strong CAM photosynthesis would rely on the differential expression of circadian clock genes during the late afternoon, and in a larger and more elaborated gene regulatory network. In addition, our results reveal the importance of the classic abiotic stress response associated to ABA into promote the shift between CAM intensity and C3 photosynthesis. These results indicate that the transition from C3 into CAM, even in plants that have evolved to do so, requires an important gene expression rewiring, making the introgression of CAM into C3 crop plants a not so straightforward process.

## Acknowledgments and Funding

We specially thanks to A. Miquel, JP Parra-Rojas, for their valuable help in the lab and the field work; to A. Miyasaka and D. Andrade for physiological instrumental facilitation and technical support; to all members of the MUCILAB, especially S. Saez-Aguayo and A. Largo-Gossens for their constant support, comments, and ideas. This work was funded by FONDAP Center for Genome Regulation (CRG) 15090007 to A.O.; ANID-Fondecyt Potsdoctorado 3510588 to P.O.; ANID-Fondecyt Iniciación 11150107 to R.N-P.; ANID-Fondecyt Iniciación 11171175 to A.M; ANID-Fondecyt regular 1200804 to C.M.

The authors declare no competing financial interests.

## Author contributions

P.O, C.M, A.M and A.O designed the study and analyses. P.O, D.O and M.T performed the field collection and ecophysiological analyses. T.C and A.R assembled and quantified the transcriptome. P.O, A.M and R.N-P performed data analysis. P.O, R.N-P performed bioinformatics and statistical data analysis. A.O and C.M provided research opportunity and funding. P.O, A.M, A.O and R.N-P wrote the manuscript.

## Supplemental Material

**Fig. S1. Analysis of soil type at the selected samples collection sites.** The HWSD Viewer from the Harmonized World Soil Database (version 1.2) was used to localize the sites the selected samples collection sites and determine the type of soil they were associated to. Arenosols are sandy soils featuring very weak or no soil development; Calcisols are soils with accumulation of secondary calcium carbonates and Regosols are soils with very limited soil development.

**Fig. S2. Statistical assessment of the differences between Sites S1, S2 and S3 plants ecophysiological parameters.** Differences among sites S1, S2 and S3 plants, in terms of nocturnal acid accumulation (Nocturnal_leaf_acid), isotopic carbon ratio (δ^13^C), leaf mass per area (LMA); succulence (SWC); total Chlorophyll/Carotenoids (ratio_Ctot:Car); Carbon to Nitrogen ratio (ratio_C:N) and photosynthetic pigments ratio (ratio_Chla:Chlb), were assessed by means of ANOVA, for all analysis but SWC. For SWC, given the data did not displayed homogeneity among samples, a Kruskal Wallis statistical test was performed. N = 10 (A). A PCA was also performed using these samples (B).

**Fig. S3. GO data clustering.** GO terms acquired from REVIGO were further summarized using the mclust method available at the simplifyEnrichment R/Bioconductor package version 1.2.0.The eight clusters that summarize the 64 GO terms are shown at the right side of the graph together with a similarity degree scale.

**Fig. S4. Normalized expression profiles from differentially expressed genes from S1 and S3 sites plants, at dawn and dusk.** Gene expression profiles associated to each community (topological module) of the four GRNs assessed in this work are displayed. Gene expression profiles are defined as the normalized counts (expression) divided by the mean normalized counts across all conditions. S1D - S1 plants genes with higher expression at dawn; S1N - S1 plants genes with higher expression at dusk; S3D - S3 plants genes with higher expression at dawn; S3N - S3 plants genes with higher expression at dusk.

**Table S1:** Data of putative CAM key gene expression regulators.

**Table S2:** qPCR primers description.

**Table S3:** *Cistanthe longiscapa* genes that were functionally annotated.

**Table S4:** Genes with a differential expression between dawn and dusk, for the S1 and S3 sites.

**Table S5:** Results from the GO and pathways analysis of the genes with a differential expression between dawn and dusk at the sites S1 and S3.

**Table S6:** Data of circadian clock genes assessed in this work.

**Supplementary File 1:** Filtered and normalized gene expression of the genes assessed for differential expression.

